# Time-of-day effects on motor learning

**DOI:** 10.1101/2022.10.03.510578

**Authors:** Charlène Truong, Célia Ruffino, Jérémie Gaveau, Olivier White, Pauline M. Hilt, Charalambos Papaxanthis

## Abstract

While the time-of-day significantly impacts motor performance, its effect on motor learning has not yet been elucidated. Here, we investigated the influence of the time-of-day on skill acquisition (i.e., skill improvement immediately after a training-session) and consolidation (i.e., skill retention after a time interval). Three groups were trained at 10 a.m. (G10_am_), 3 p.m. (G3_pm_), or 8 p.m. (G8_pm_) on a finger-tapping task. We recorded the skill (i.e. the ratio between movement duration and accuracy), before and immediately after the training to evaluate skill acquisition, and after 24 hours, to measure skill consolidation. We did not observe any difference in acquisition according to the time of the day. However, we found an improvement in performance 24 hours after the evening training (G8_pm_) while the morning (G10_am_) and the afternoon (G3_pm_) groups deteriorated and stabilized their performance, respectively. Furthermore, two control experiments (G8_wake_ and G8_sleep_) supported the idea that a night of sleep contributes to the skill consolidation of the evening group. These results show an influence of time-of-day on the consolidation process, with better consolidation when the training is carried out in the evening. This finding may have an important impact on the planning of training programs in sports, clinical, or experimental domains.

## Introduction

Motor learning is essential for the development of new motor skills and the perfection or preservation of the existing ones (Krakauer et al., 2019). Fundamental to the acquisition of a profuse motor repertoire is practice. The positive effects of practice on skill acquisition can be observed with different timescales, from a single session to several days, weeks, or months (Dayan and Cohen, 2011). Typically, a single practice session leads to a rapid increase in accuracy and/or speed until the reach of an asymptotic level of performance (Doyon and Benali, 2005). This fast learning process, defined as *motor acquisition*, is the first step leading to the formation of new motor memories. Within additional practice sessions or between them (i.e., during the rest period following practice, defined as off-line learning), a consolidation process transforms the new initially labile motor memory into a robust motor memory (Robertson et al., 2004a; Ruffino et al., 2021). Notably, several studies, including diurnal and nocturnal sleep groups, have highlighted the key role of sleep on motor skill consolidation (Fischer et al., 2002; Walker, 2003; Rickard et al., 2008; Debas et al., 2010; Fogel et al., 2014). Both acquisition and consolidation processes are associated with neuronal adaptations and plastic modulations at several levels of the central nervous system (for review see Dayan and Cohen, 2011).

Optimization of skill learning is important for many professions and the literature on this topic is abundant. The amount of training (Driskell et al., 1992; Karni et al., 1995), the distribution of rest periods during a training session (Rickard et al., 2008; Verhoeven and Newell, 2018), or the variation of skills within a single training session (Shea and Morgan, 1979; Neville and Trempe, 2017) are some examples. Astonishingly, the optimal time-of-day for motor learning has not yet retained great attention, even though several studies have shown that motor and mental performances fluctuate across the day, following a circadian basis (~24 h) (Atkinson and Reilly, 1996; Drust et al., 2005; Gueugneau and Papaxanthis, 2010). For example, daily variations have been observed for maximal voluntary contractions (Guette et al., 2005), spontaneous motor tempo (Hammerschmidt and Wöllner, 2022), speed/accuracy tradeoff of actual and mental movements (Gueugneau et al., 2017), handwriting (Jasper et al., 2009), and tennis service (Atkinson and Speirs, 1998). In all studies, better performances have been consistently reported in the late afternoon than early in the morning.

Sporadic and contradictory information regarding the time-of-day influence on motor learning can be extracted from existing studies, in which the main objective was the role of sleep in motor learning. Precisely, some investigations did not find differences in skill learning between morning and evening training (Korman et al., 2003; Blischke et al., 2008; Kvint et al., 2011; Sale et al., 2013). Others tempered this finding by showing that the time-of-day positively affects the expression of motor learning (Keisler et al., 2007; Holz et al., 2012; Truong et al., 2022). Up to now, the daily variation in motor learning appears to be an unresolved issue. Consequently, it is motivating not only to investigate if there is an optimal period during the day to practice and learn a motor skill but also to elucidate if this optimal period is equally auspicious for motor acquisition and/or consolidation.

Here, we designed a specific experimental protocol to explore the influence of time-of-day in skill acquisition and consolidation. We firstly composed three groups that practiced at different times-of-day (Day 1): 10 a.m. (G10_am_), 3 p.m. (G3_pm_), and 8 p.m. (G8_pm_). In the first session, participants were trained on a finger tapping task to measure skill acquisition, namely the enhancement in skill performance during and immediately after practice. In the second session, scheduled 24 hours after the first session (Day 2), we measured skill consolidation, that is, the amount of skill retention or improvement. Following the time-of-day literature (Atkinson and Reilly, 1996), we hypothesized that skill acquisition and consolidation should be better in the afternoon compared to the morning. We also motivated this premise by neurophysiological findings showing that physiological mechanisms are modulated throughout the day, such as the cortisol diurnal secretion related to LTP-like plasticity in the motor cortex (Sale et al., 2008) and the degree of hippocampus activation (Dolfen et al., 2019), associated with consolidation (Albouy et al., 2015; King et al., 2021).

When testing motor memory consolidation among different training periods during the day, one is faced with a double problem: the passage of time and sleep, since both contribute to motor consolidation (Robertson et al., 2004b; Cohen et al., 2005). For that reason, we included two supplemental groups, both trained in the evening (8 p.m., Day 1). The G8_sleep_ was tested 14 hours after the practice session (at 10 a.m. on Day 2). By comparing the G8_sleep_ with the G8_pm_, we controlled for the time of being awake the Day 2 (2 hours for the first, 12 hours for the second). By comparing the G8_sleep_ with the G10_am_, we controlled for the specific time of testing skill consolidation on Day 2 (10 a.m. for both), whatever the training period on Day 1. The G8_wake_ was tested 2 hours after the practice session to test the positive effect, if any, of the passage of time before the night of sleep. If only the night of sleep contributes to the development of off-line learning, we should not observe change before the night of sleep, while offline gains should be observed immediately after the night of sleep.

## Method

### Participants

Fifty-nine healthy adults participated in the current study after giving their informed consent. All were right-handed (mean score 0.8 ± 0.2), as defined by the Edinburgh handedness questionnaire (Oldfield, 1971), and free from neurological or physical disorders. Due to the nature of the motor task (finger typing) used in the present study, we excluded musicians and professional typists. Participants were randomly assigned into five groups: the G10_am_ (n = 12, 8 females, mean age: 25 ± 6 years old), the G3_pm_ (n = 12, 7 females, mean age: 24 ± 6 years old), the G8_pm_ (n = 12, 6 females, mean age: 23 ± 2 years old), the G8_sleep_ (n = 12, 5 females, mean age: 23 ± 4 years old), and the G8_wake_ (n = 11, 4 females, mean age: 26 ± 3 years old). Data from participants of G10_am_ and G3_pm_ was reused from a previously published study (Truong et al., 2022). The experimental design was approved by the regional ethic committee (Comité de Protection des Personnes - Région EST) and conformed to the standards set by the Declaration of Helsinki.

All participants were requested to be drug- and alcohol-free, to not change their habitual daily activities (e.g., cooking, computer use, handiwork), and to not make intensive physical activity during the 24 hours preceding the experiment. They were all synchronized with a normal diurnal activity (8 a.m. ± 1 hour to 12 a.m. ± 1 hour alternating with the night). We examined the chronotype of each participant using the Morningness-Eveningness Questionnaire (Horne and Ostberg, 1976). In this test, scores range from 16 to 86 and are divided into five categories: evening type (score 16 to 30, n=2 for our study), moderate evening type (score 31 to 41, n=5), intermediate type (score 42 to 58, n=45), moderate morning type (score 59 to 69, n=6), and morning type (score 70 to 86, n=1). There were no significant differences between groups regarding the chronotype (on average; G10_am_: 51 ± 11; G3_pm_: 50 ± 9; G8_pm_: 52 ± 10; G8_wake_: 54 ± 10; G8_sleep_: 47 ± 8; one-way ANOVA F_4,54_ = 1.05, *p* = 0.39). We verified the sleep quality of each participant with the Pittsburgh Sleep Quality Index (Buysse et al., 1989). The general score in this questionnaire ranges from 0 (no particular difficulties sleeping) to 21 (major difficulties sleeping). The results indicated good sleep quality for all groups (grand average; G10_am_: 5 ± 1, G3_pm_: 4 ± 1, G8_pm_: 5 ± 3; G8_wake_: 4 ± 3; G8_sleep_: 5 ± 3; oneway ANOVA: F_4,54_ = 0.28, *p* = 0.89).

### Experimental device and procedure

Participants were comfortably seated on a chair in front of a keyboard. We employed a computerized version of the sequential finger tapping task (see Truong et al., 2022). Specifically, participants were requested to tap as accurately and fast as possible with their non-dominant hand the following sequence: 1-4-2-3-1-0. Each key was affected by a specific finger: 0-thumb, 1-index, 2-middle, 3-ring, and 4-little. One trial was composed of six sequences. At the beginning of each trial, participants pressed the key ‘0’ with their thumb to start the chronometer and they accomplished the 6 sequences continuously. Pressing the key ‘0’ at the end of the 6^th^ sequence stopped the chronometer and ended the trial (Fig. 1A).

**Figure 1.**
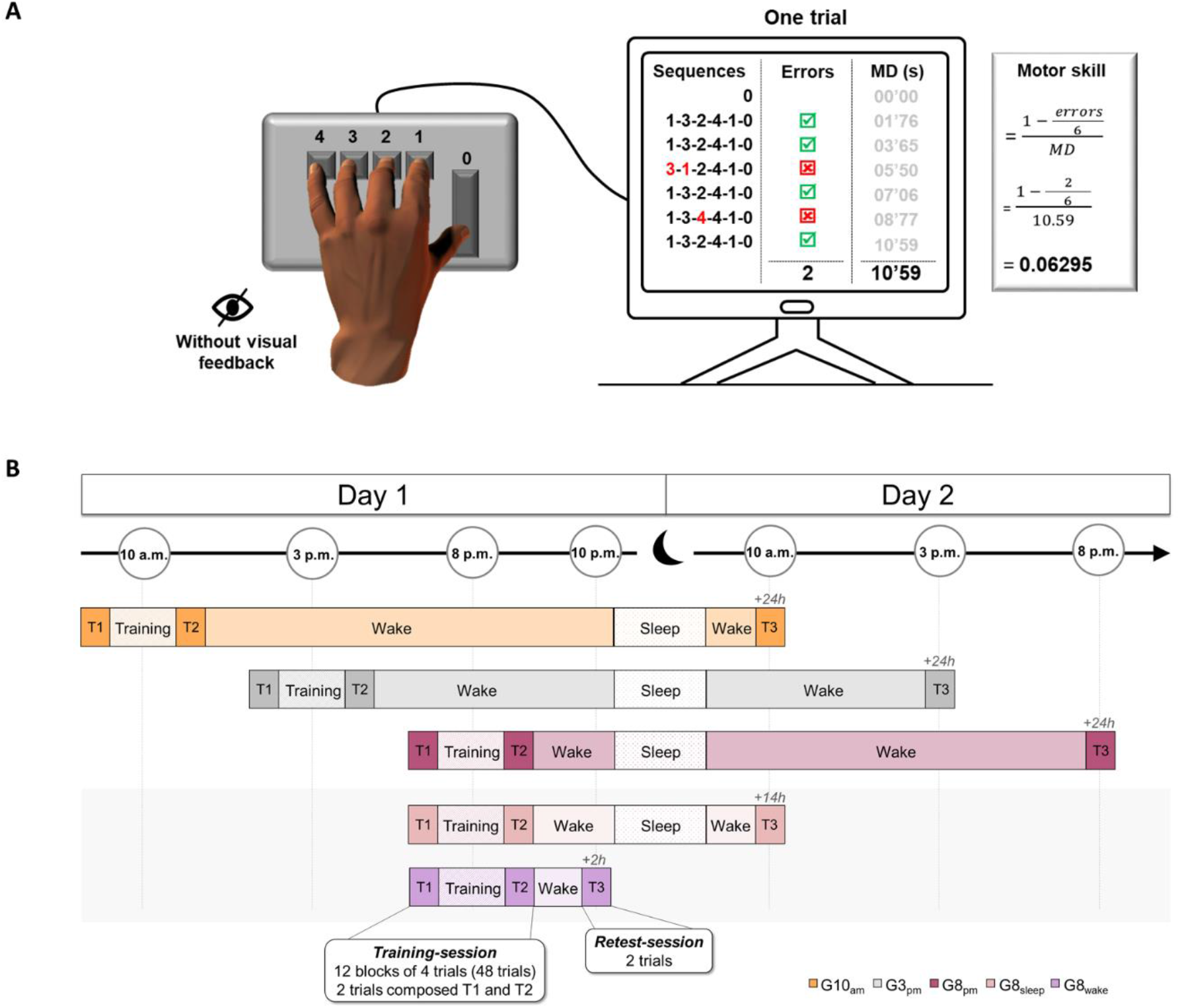
Experimental design. **(A)** Participants’ hand position on the keyboard and the computerized version of the sequential finger-tapping task. Each key was affected by a specific finger: 0-thumb, 1-index, 2-middle, 3-ring, and 4-little. One trial included 6 successive sequences: 1–4–2–3–1–0. One trial was composed of six sequences. At the beginning of each trial, participants pressed the key O’ with their thumb to start the chronometer and they accomplished the 6 sequences continuously. Pressing the key ‘0’ at the end of the 6^th^ sequence stopped the chronometer and ended the trial. We recorded the number of false sequences and the duration of the whole trial (MD). **(B)** Experimental procedure. On Day 1, participants performed a training-session at 10 a.m. (G10_am_), 3 p.m. (G3_pm_), and 8 p.m. (G8_pm_, G8_sleep_, G8_wake_). All participants carried out 48 trials (12 blocks of 4 trials, with 5-s rest between trials and 30-s rest between blocks). The pre-test (Tl) and the post-test (T2) were composed, respectively, of the first two trials (1 and 2) and the last two trials (47 and 48). Then, for the retest-session, all participants accomplished two supplementary trials (T3). The G10_am_, G3_pm_ and G8_pm_ were tested 24 hours after the training (Day 2). The G8_sleep_ and the G8_wake_ were tested 14 hours after the training (Day 2 at 10 a.m.) and 2 hours after the training (Day 1 at 10 p.m.), respectively.

The experiments included two *sessions*, scheduled on two consecutive days (Day 1 and Day 2), except for the G8_wake_ (tested only on Day 1, see Fig. 1B). On Day 1, participants were tested at 10 a.m. (G10_am_), 3 p.m. (G3_pm_) or 8 p.m. (G8_pm_, G8_wake_, and G8_sleep_). They carried out 48 trials (12 blocks of 4 trials, with 5-s rest between trials and 30-s rest between blocks). The score in the first two trials (1 and 2) and the last two trials (47 and 48) composed the pre-test (T1) and the post-test (T2), respectively. The remaining trials (n = 44; from the 3^rd^ to the 46^th^ trial) constituted the training trials. To familiarize themselves with the protocol, all participants accomplished two trials at a natural speed. For the *retest-session*, all participants carried out two trials (T3). The G10_am_, G3_pm_, and G8_pm_ were re-tested 24 hours (Day 2) after the first *training-session*. The G8_sleep_ was re-tested14 hours (Day 2 at 10 a.m.) after the first *trainingsession*. The G8_wake_ was re-tested 2 hours (Day 1 at 10 p.m.) after the first *training-session*.

The vision of the non-dominant hand was hidden using a box during the whole protocol. The sequence’s order, however, was displayed on the box and thus visible to the participants during the whole experiment. Note that no information concerning motor performance (i.e., time or typing errors) was provided to the participants.

### Data recording and analysis

For each trial, a Visual Basic for Applications program (Microsoft, Excel) recorded movement accuracy and duration. The accuracy (‘Error rate’) was defined as the number of false sequences throughout one trial (0 = no false trial; 6 = all trials false). If the participants made one or more mistakes in one of the sequences, this sequence was counted as false (see Fig. 1A). Error rate was defined as the percentage of the number of errors during a trial:

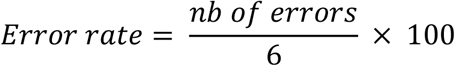

Movement duration (second) was defined as the time interval between the start of the trial (when the participants pressed the key ‘0’) and the end of the trial (when the participants pressed the key ‘0’ at the end of the 6^th^ sequence).

These two parameters (Movement duration and Error rate) are related by the speed-accuracy tradeoff function (Shmuelof and Krakauer 2012). Ascertaining, thus, that motor skill (i.e., the training-related change in the speed-accuracy trade-off function) has been improved is not possible when duration and accuracy change in opposite directions. For that reason, we compute a composite ratio to describe motor skill as follows:

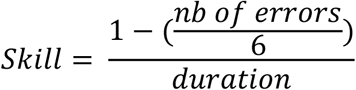

In that formula, skill increases when the ratio increases.

Gains between sessions were calculated as follows:

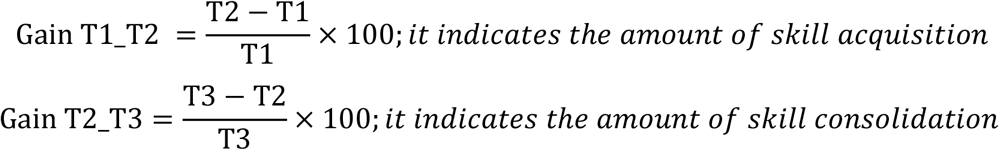

### Statistical analysis

For all data, we verified the normality and sphericity using Shapiro–Wilk test and Mauchly’s test, respectively. For all statistical analyses, the significance level was fixed at 0.05 and the observed power was superior to 0.8.

First, we compared the G10_am_, G3_pm_, and G8_pm_. For skill and movement duration, we applied repeated measures (rm) ANOVA with *group* as between-subjects factor (G10_am_, G3_pm_, and G8_pm_) and *session* as the within-subjects factor (T1, T2, and T3). Gains in motor skill (T1_T2 and T2_T3) were analyzed by one-way ANOVA between groups. When necessary, the *post-hoc* analyses were performed by applying Newman-Keuls tests. Moreover, each gain was compared with the reference value *‘zero’* (0).

As the error rate did not follow a normal distribution (Shapiro-Wilk test, p < 0.05), we used two-tailed permutation tests (5000 permutations; MATLAB function mult_comp_perm_t1). P-values were corrected for multiple comparisons using the Benjamini-Hochberg False Discovery Rate (MATLAB function fdr_bh).

In addition, the trial-by-trial (n=48) evolution of individual skill performance during the whole training session was analyzed by a power-law function:

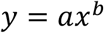

where *y* is the skill performance, *x* is the trial number, *a* is the learning amplitude, and the exponential *b* is the learning rate. We used one-way ANOVA between groups for the learning rate (b). To further analyze the form of each learning curve (represented by the powerlaw function), we determined the moment at which the skill performance tends to stabilize. Precisely, we computed the derivative of the fitted power function for each participant. To define the moment at which skill progression tends to be null, we computed the time point at which the derivative reaches 0.0001 above its minimum value for each participant. We compared these values between groups using a one-way ANOVA. Since our threshold was arbitrarily determined, we performed, as a control, the same analysis with threshold values of 0.0002, 0.0003, 0.0004, and 0.0005 (results of these analyses are presented in Supplementary result 1).

To test whether an eventual improvement in skill performance at T3 was due to offline learning rather to the continuation of training (see Pan and Rickard, 2015), we extrapolated the individual skill performance during the 48 trials to an additional two trials (49 and 50) in G8_pm_. We determined the gain T2_T3_pred_ of this predicted evolution of training (T3_pred_) and compared it with the gain T2_T3 of G8_pm_ by independent samples T-test.

We also tested skill and movement duration of G8_sleep_ using rmANOVA with *session* as within-subject factor (T1, T2, and T3) and performed Newman-Keuls post hoc. As in the first analysis, the error rate was analyzed by permutation tests, corrected by the Benjamini-Hochberg False Discovery Rate, and the skill gains (T1_T2 and T2_T3) of G8_sleep_ were compared to the reference value *‘zero’* (0). In addition, we compared skill gains of G8_sleep_ with G8_pm_ with independent t-tests. Finally, we performed analogous statistical analyses for the G8_wake_.

## Results

Figure 2A shows the average values (+SD) of skill for G10_am_, G3_pm_, and G8_pm_. rmANOVA revealed a significant interaction effect (*group* x *session;* F_4,66_ = 5.67, p < 0.001, η^2^ = 0.26). *The post-hoc* analysis did not show significant differences between groups in T1 (in all, p > 0.7). A separate analysis for speed (rm ANOVA: F_4,66_ = 2.61, p < 0.05, η^2^ = 0.14; *post-hoc* analysis: in all, p > 0.58) and accuracy (in all, T < 0.35, p > 0.97) gave similar results (see Table 1). Skill significantly enhanced after training (T2) for all groups (T1 versus T2 *post-hoc* comparisons: in all, p < 0.001; see Fig 2A). Note that a distinct analysis for speed and accuracy showed that skill improvement between T1 and T2 was mainly due to faster movements (in all, p < 0.001; see Table 1) rather than more accurate movements (in all, T < 1.99, p > 0.12; see Table 1). Figure 2B illustrates the average (+SD) acquisition gains (T1_T2) in skill. The comparison of T1_T2 gain with the reference value *zero (0)* showed significant improvement for all groups (in all, t > 5.87, p < 0.001). It is of interest that skill acquisition was not significantly different between groups (one-way ANOVA: F_2,33_ = 0.41, p = 0.67, η^2^ = 0.02). To further explore skill acquisition, we examined the learning curves of skill (Fig 2C). The analysis did not reveal a significant difference between groups (factor b: F_2,33_ = 0.54, p = 0.54, η^2^ = 0.04), indicating that the speed of skill improvement did not differ according to the time-of-day. Additionally, we did not find any modulation of the moment at which the skill enhancement tended to stabilize (G10_am_: 26 ± 5 trials; G3_pm_: 27 ± 3 trials; G8_pm_: 27 ± 3 trials; F_2,33_ = 0.37, p = 0.69, η^2^ = 0.02), whatever the threshold chosen (see also Supplementary results 1). Overall, the above results indicate that skill improvement during the training sessions does not differ between groups, suggesting that the acquisition process is independent of the time-of-day.

**Figure 2.**
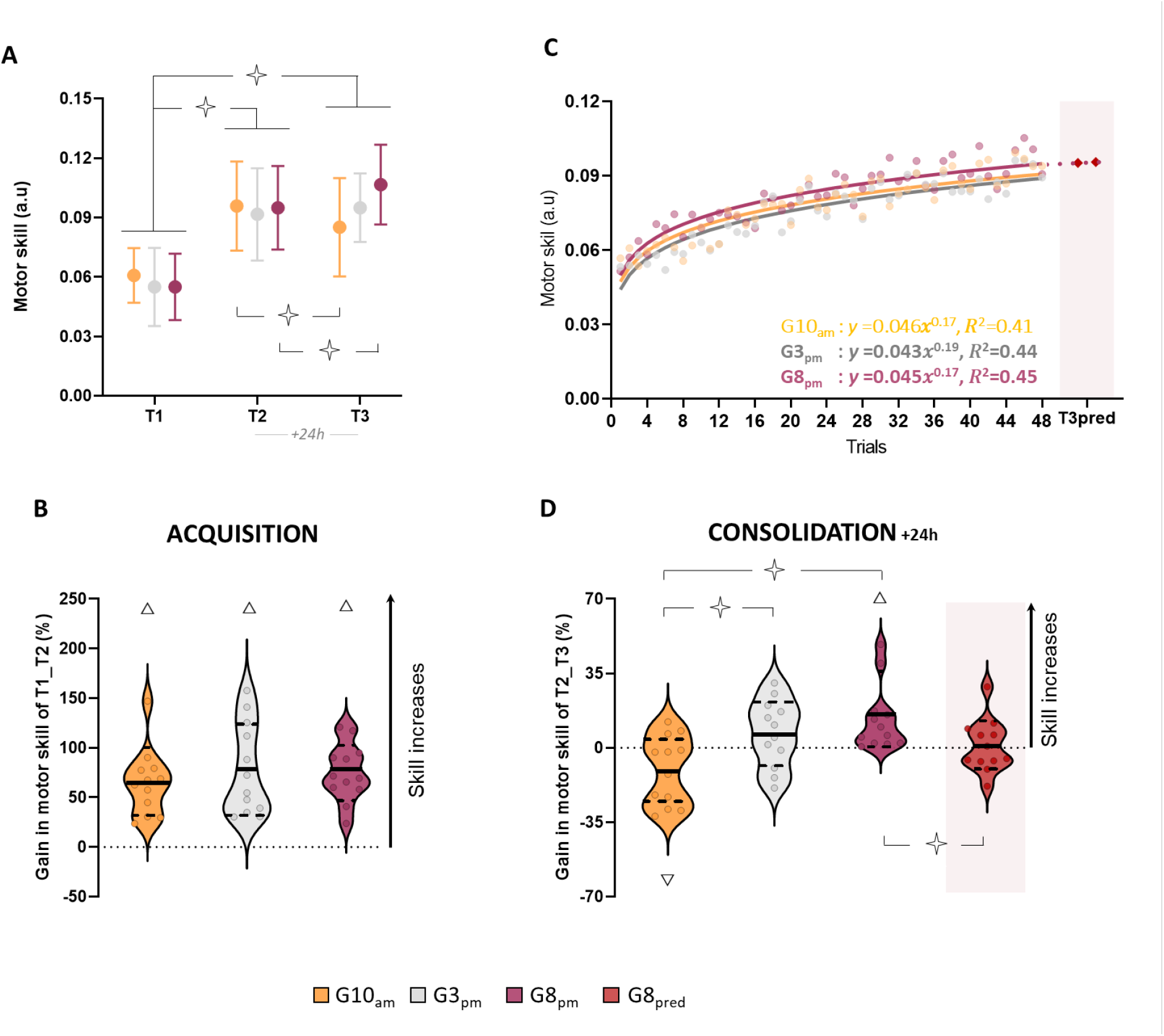
Skill performance for the G10_am_, G3_pm_, and G8_pm_ groups. **(A)** Average values and standard deviations (+SD) of skill in T1, T2, and T3 for each group. **(B)** Violin plots for the percentage of acquisition gain in skill (Tl_T2). Thick and thin horizontal lines mark mean and SD, respectively. Dots represent individual data per condition. **(C)** Trial-by-trial plotting of skill evaluation during the training and the corresponding power law functions. Diamonds are the extrapolation to the two additional trials composing the T3*_pred_*. **(D)** Violin plots for the percentage of consolidation gain in skill (T2_T3) for each group and the G8_pred_ (prediction of gain by extrapolation for G8_pm_). Thick and thin horizontal lines mark mean and SD, respectively. Dots represent individual data per condition. Open stars indicate significant differences between groups or sessions. Triangles indicate significant differences from the value *zero*.

**Table 1.**
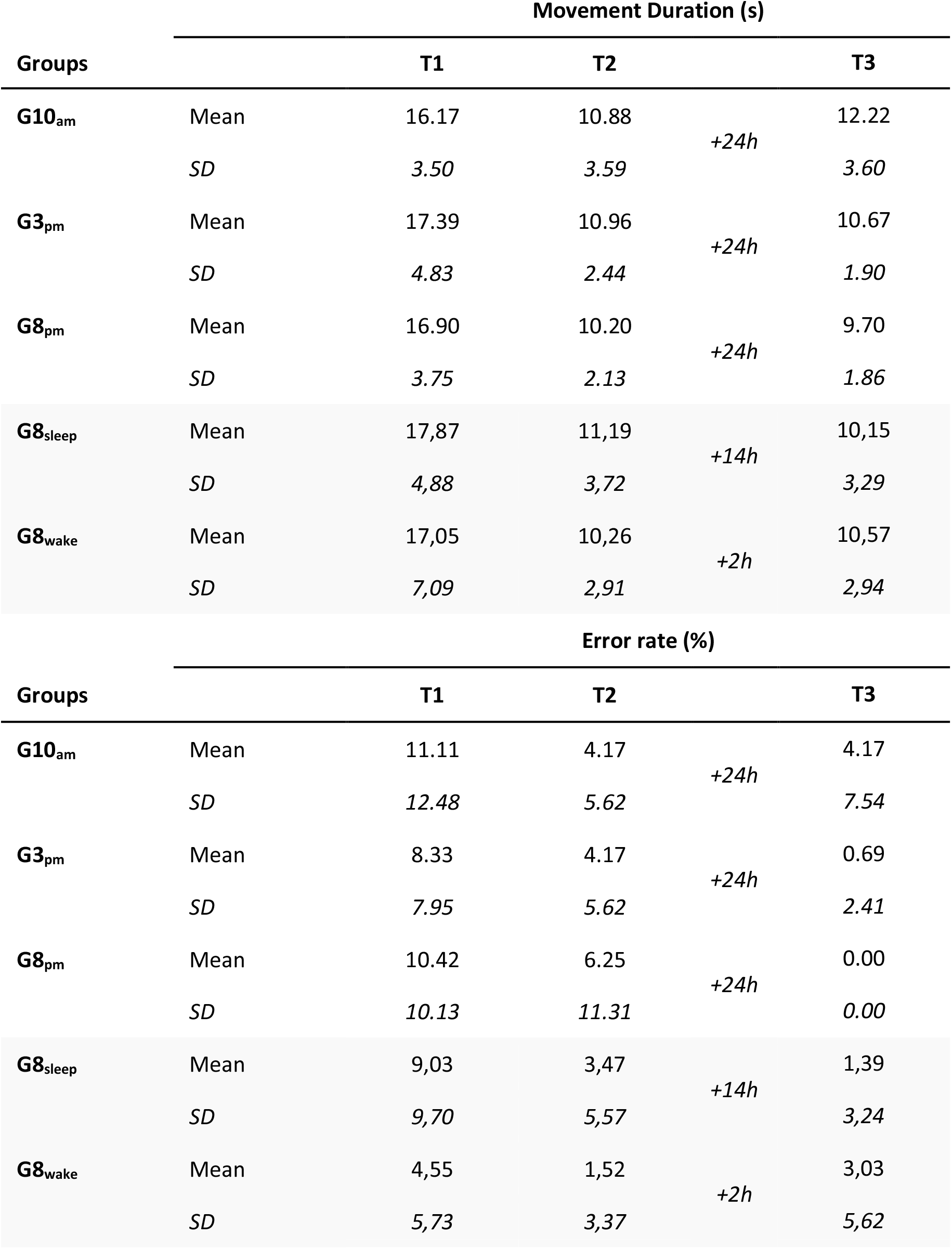
Average value (+SD) of Movement duration (s) and Error rate (%) for the T1, T2, and T3 and each group.

Regarding skill consolidation, we found notable differences between groups, explaining the interaction effect found for skill (*group* x *session*). Precisely, one day after training (T2 versus T3; *post-hoc* analysis), there was a deterioration in skill performance for the G10_pm_ (p = 0.03), a stabilization for the G3_pm_ (p = 0.71), and further improvement for the G8_pm_ (p = 0.02). A separate analysis did not reveal statistical differences between T2 and T3 neither for speed (in all, p > 0.20) nor for accuracy (in all, T < 1.94, p > 0.09), suggesting that changes in skill consolidation were due to small changes in both speed and accuracy (see Table 1). Figure 2D illustrates the average (+SD) consolidation gains (T2_T3) in skill. The comparison of T2_T3 gain with the reference value zero (0) showed a deterioration for the G10_am_ (t = −2.27, p = 0.04; 9/12 participants decreased their performance), a stabilization for the G3_pm_ (t = 1.49, p = 0.17; 8/12 participants increased their performance), and an improvement of skill for the G8_pm_ (t = 3.02, p = 0.01; 12/12 participants increased their performance). One-way ANOVA (F_2,33_ = 7.44, p = 0.002, η^2^ = 0.32) showed that T2_T3 gain of the G10_am_ significantly differed from that of the G3_pm_ and G8_pm_ (for both, p < 0.05).

Importantly, skill improvement for the G8_pm_ at T3 was not due to a simple continuation of the practice, namely to the improvement that this group would obtain if two more trials would be carried out on Day 1 (50 trials instead of 48), but rather to offline learning. Indeed, the T2_T3 gain of G8_pm_ was significantly different from the T2_T3_pred_ gain (t = 2.21, p < 0.05; see Fig. 2C and 2D). Despite group differences in offline skill improvement after training (T2 versus T3), all groups acquired better skill performance one-day later (T3) compared to their initial performance (T1 versus T3; in all, p < 0.001).

Note that to exclude any effects of different chronotypes, we conducted the same statistical analyses without the extreme and moderate chronotypes and we obtained the same results (see Supplementary results 2).

Overall, the consolidation process (i.e., skill measured 24h later) was affected by the time of the day wherein the training was carried out. This finding suggests that physical training scheduled close to the night of sleep may enable offline learning, while that scheduled in the morning may lead a forgetting. Before asserting such a premise, however, we controlled for alternative issues.

### Control Experiment 1

In the main experiment, the groups were retested (T3) at different times the Day 2 to respect the 24h delay. Precisely, the participants of the G8_pm_ were retested twelve hours after waking up (8 a.m. – 8 p.m.), compared to those of the other groups who were tested 2 hours (G10_am_; 8 a.m. – 10 a.m.) or 7 hours (G3_pm_; 8 a.m. – 3 p.m.) after waking up. Therefore, one must exclude the possible effects of the time being awake the day after training (Day 2), which could positively contribute to skill consolidation. To evaluate such an effect, the G8_sleep_ performed the training at 8 p.m. and was retested the next day at 10 a.m. (i.e., a total of 14 hours after the training but 2 hours after waking up; see Fig 1B). If offline learning is due to training close to the night of sleep and not to the time of being awake the day after, the individuals of the G8_sleep_ should show comparable improvement with those of the G8_pm_.

rmANOVA revealed a significant effect of *session* for skill (F_2,22_ = 65.25, p < 0.001, η^2^ = 0.85; see Fig 3A). The *post-hoc* analysis showed significant differences between all measurements (T1-T3; p < 0.05). After training, skill acquisition (T1_T2 gain) significantly improved (compared to reference value *zero* (0); t = 6.07, p < 0.001; Fig. 3B). No significant difference between the G8_sleep_ and the G8_pm_ was found for the gain T1_T2 (t= 0.07, p = 0.94). After the night of sleep (T2_T3), we observed a significant offline improvement (compared to reference value *zero* (0): t = 4.44, p < 0.001; 10/12 participants increased their performance). This enhancement was not different (t = −0.18, p = 0.86) to that obtain for the G8_pm_. In addition, T2_T3 gain in skill for the G8_sleep_ was significantly greater than for the group G10_am_ (t= 3.80, p < 0.001). Note that speed and accuracy of G8_sleep_ (see Table 1) contributed in the same way as G8_pm_ for the skill acquisition, i.e., improvement in speed (T1 versus T2; F_2,22_ = 66.87, p < 0.001, η^2^ = 0.86; *post hoc* analysis: p < 0.001) but not in accuracy (T = 1.72, p > 0.21), and consolidation, i.e., slight improvement, although significant, for both speed and accuracy (T2 versus T3; for movement duration: *post hoc* analysis: p > 0.16; for error rate: T = 1.12, p > 0.57).

**Figure 3.**
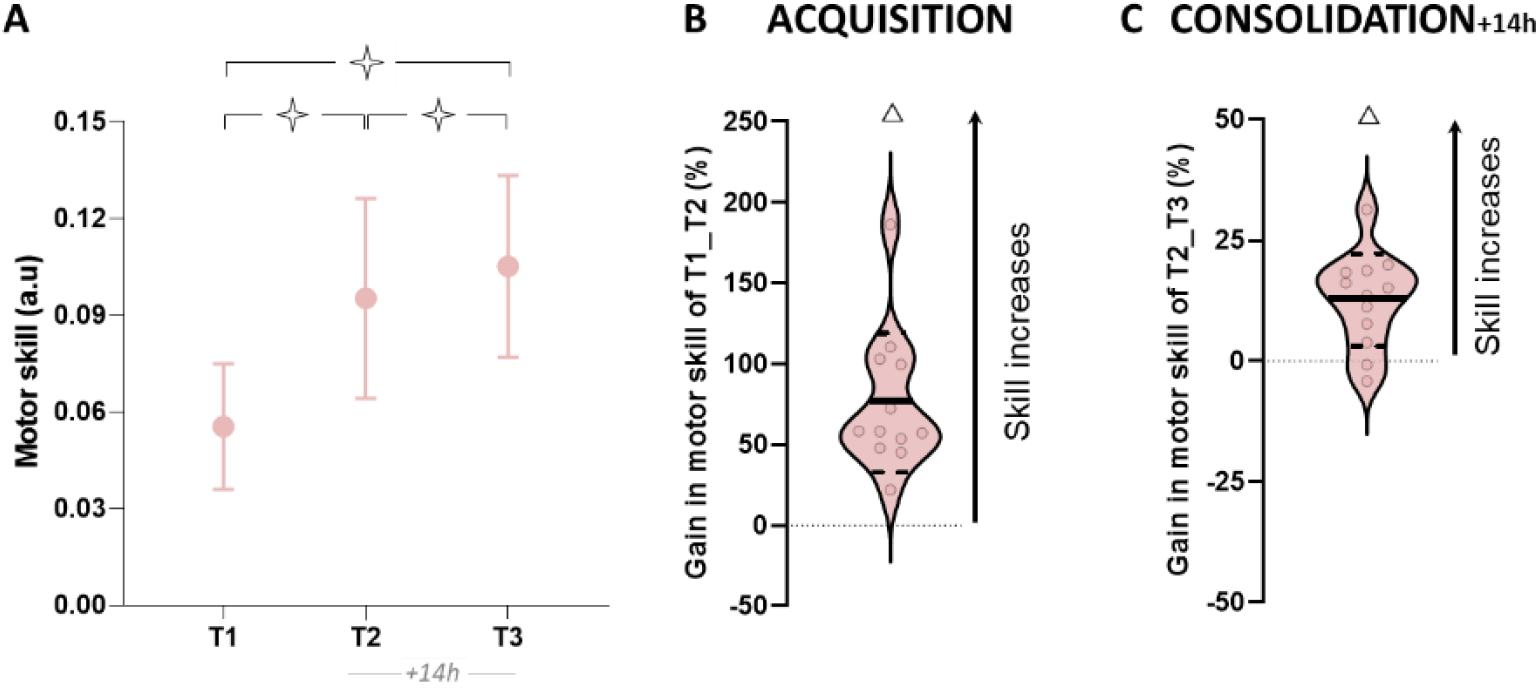
Skill performance for the G8_sleep_ group **(A)** Average values (+SD) of skill in T1, T2, and T3. Open stars indicate significant differences between sessions. **(B)** Violin plots for the percentage of acquisition gain in skill (T1_T2). **(C)** Violin plots for the percentage of consolidation gain in skill (T2_T3). Thick and thin horizontal lines mark mean and SD, respectively. Dots represent individual data per condition. Triangles indicate significant differences from the value *zero*.

According to these results, skill consolidation seems to be independent of the time being awake the day after the *training-session*, further reinforcing the premise that sleep significantly contributes to skill consolidation.

### Control Experiment 2

To directly attribute the skill consolidation to sleep, one must also examine for any positive effect of the passage of time between training and sleep. Approximately, 4h elapsed between the beginning of training (8 p.m.) and time of sleep (12 a.m.) for the G8_pm_, and therefore, one could not exclude any improvement/consolidation in skill performance during this period. To evaluate such an effect, the G8_wake_ carried out the training at 8 p.m. and was retested 2 hours later at 10 p.m.

Figure 4A illustrates the average values (+SD) of skill for G8_wake_. rmANOVA revealed a significant session effect (F_2,20_ = 25.56, p < 0.001, η^2^ = 0.72). The *post-hoc* analysis showed a significant difference between the first session (T1) and the other sessions (T2 and T3; for both, p < 0.001), while no significant difference was detected between T2 and T3 (p = 0.44). After training, skill acquisition (T1_T2 gain) significantly improved (compared to reference value *zero* (0): t = 4.34, p = 0.001; see Fig 4B). Note that, T1_T2 gain was not significantly different to G8_pm_ (t = −0.16, p = 0.86). After 2 hours, T2_T3 gain in skill did not significantly improve (compared to reference value *zero* (0): t = −1.07, p = 0.31; see Fig 4C). In addition, the T2_T3 gain in skill for this group was significantly inferior to the offline gain of the G8_pm_ (t = −2.99, p < 0.01). Like the previous group, speed and accuracy of G8_wake_ (see Table 1) contributed in the same way as G8_pm_ for the skill acquisition (T1 versus T2; significant improvement for speed: F_2,20_ = 15.23, p < 0.001, η^2^ = 0.60; *post hoc* analysis: p < 0.001; no significant improvement for accuracy: T = 1.51, p > 0.53). However, for the consolidation, we observed slight deterioration, but no significant, of both speed (T2 versus T3; *post hoc* analysis: p > 0.82) and accuracy (T = 0.77, p > 0.66).

**Figure 4.**
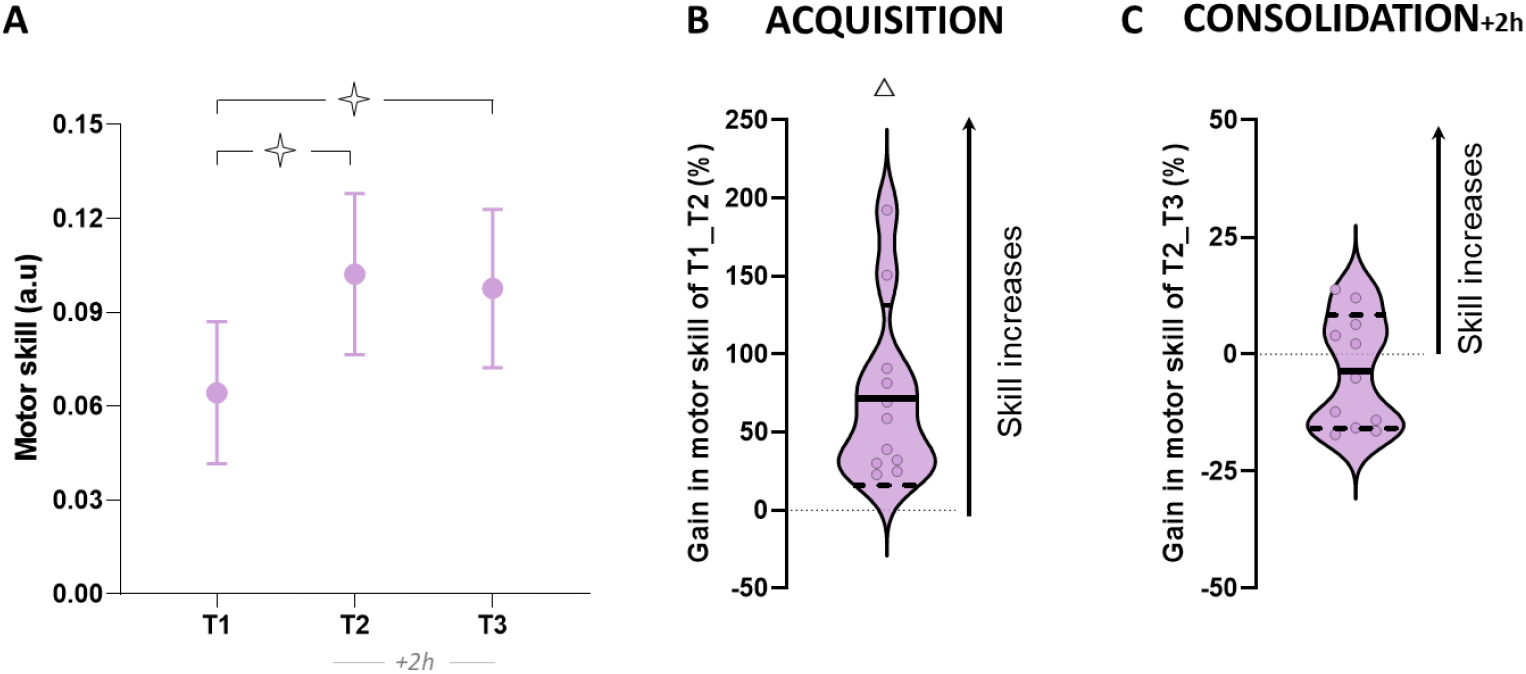
Skill performance for the G8_wake_ group **(A)** Average values (+SD) of skill in T1, T2, and T3. Open stars indicate significant differences between sessions. **(B)** Violin plots for the percentage of acquisition gain in skill (T1_T2). **(C)** Violin plots for the percentage of consolidation gain in skill (T2_T3). Thick and thin horizontal lines mark mean and SD, respectively. Dots represent individual data per condition. Triangle indicates significant differences from the value *zero*.

Overall, these results support the contribution of sleep in the offline improvement in skill performance when the training is planned close to sleep.

## Discussion

In the current study, we examined the influence of the time-of-day on motor skill acquisition and consolidation following physical practice on a finger tapping task. Our findings did not show any effect of the time-of day (10 a.m., 3 p.m., and 8 p.m.) in the acquisition process (i.e., skill improvement during the training session) as all groups equally improved their skill performance. Skill consolidation, however, significantly differed accordingly to the time-of-day. Precisely, we observed a deterioration in skill performance 24 hours after training in the morning (G10_am_), a stabilization after training in the afternoon (G3_pm_), and an improvement after training in the evening (G8_pm_). In addition, the findings from the two control groups showed that offline gains in skill performance for the evening group were mainly due to the night of sleep than to the awakening time between training and sleep or the amount of time passed awake after sleep (G8_wake_ and G8_sleep_, respectively).

### The time-of-day effects on skill acquisition

As in many previous findings (Fischer et al., 2002; Walker, 2003; Blischke et al., 2008; Doyon et al., 2009; Kvint et al., 2011; Albouy et al., 2013; Tucker et al., 2017), all participants in our study increased skill performance following a single session of practice. Interestingly, skill improvement was comparable whatever the time-of-day in which training was provided (morning, afternoon, or evening). Furthermore, skill learning rate (see learning curves, Fig 2C) was also similar among groups and thus according to the time-of-day. These findings indicate that skill acquisition is independent of the time-of-day in which training is administrated.

These results may be surprising since clear influences of the time-of-day on behavioral and neural processes have been described (Atkinson and Reilly, 1996; Drust et al., 2005). For instance, the maximal voluntary contractions (Guette et al., 2005), speed/accuracy tradeoff of actual and mental movements (Gueugneau et al., 2017), handwriting (Jasper et al., 2009), and badminton serve accuracy (Edwards et al., 2005) fluctuate through the day. It is proposed that the naturally higher values of body temperature in the late evening compared to the morning may enhance motor performance (Teo et al., 2011; Kusumoto et al., 2021). In addition, neurophysiological mechanisms, such as Long-Term Potentiation (LTP-like plasticity) and intracortical inhibition within the primary motor cortex (M1), both engaged in skill acquisition (Ziemann et al., 2004; Rosenkranz et al., 2007; Coxon et al., 2014; Spampinato and Celnik, 2018), are also modulated by circadian rhythms. For example, Sale et al. (2008) showed that these neural mechanisms seem to vary across the day according to the cortisol hormonal circadian fluctuation. Furthermore, Lang et al. (2011) showed that the inhibitory networks on M1 (notably by GABA-B mediated), influencing neural plasticity (McDonnell et al., 2007), also fluctuated throughout the day.

In the present study, such plastic neural changes with the M1 seem to not affect skill acquisition. Possibly, plastic neural changes operating during the day may influence the consolidation rather than the acquisition of motor skills. In addition, the acquisition of motor sequences is a complex process, which is not limited to the M1 area but includes different cortico-subcortical networks such as cortico-striatal (CS) and the cortico-cerebellar (CC) networks (Doyon and Benali, 2005; Lohse et al., 2014). It is also important to note that skill acquisition includes multiple mechanisms (Smith et al., 2006; Spampinato and Celnik, 2021) and reflects the sum of different learning components, such as implicit and explicit learning (Taylor et al., 2014; Morehead et al., 2015; for review see Krakauer et al., 2019). In the present study, we did not explore these mechanisms analytically. It should be interesting to isolate these components to examine their evolution across the day to provide a better understanding of our results.

### The time-of-day effects on skill consolidation

The majority of the studies on skill consolidation showed offline learning, i.e., additional improvement of performance without practice after a night of sleep (Fischer et al., 2002; Walker et al., 2002, 2003; Korman et al., 2003; Walker, 2003; Kuriyama, 2004; Blischke et al., 2008; Doyon et al., 2009). Here, we found that this offline learning occurs only if the training takes place in the evening. More precisely, we observed different amounts of consolidation according to the training schedule, with a deterioration of motor skill for the morning group (10 a.m.), stabilization for the afternoon group (3 p.m.), and improvement for the evening group (8 p.m.) 24 hours after the training. Notably, our results showed that the offline skill improvement of the evening group was due to a consolidation process (see Fig 2C with the T3_pred_ and Fig 2D with the gain of G8_pred_), and not due to the continuation of the learning, that is to a simple relearning process (Rickard et al., 2008). In such a case, one should observe the same amount of improvement between 24 hours later (T3) and the extrapolation of learning (T3_pred_).

Additionally, we showed that offline learning did not occur with the simple passage of time just after training (G8_wake_; see control experiment 2) and that sleep is important for offline learning. We also found comparable offline improvement when the retest was scheduled the next day in the morning or the evening (G8_sleep_ vs G8_pm_; see control experiment 1), excluding the possible contribution of the retest schedule and, thus, the time being awake after the night of sleep. Many studies have reported changes in non-rapid-eye-movement (NREM) stage-2 sleep as well as reactivations of task-related memory networks during sleep following a training session (Rasch and Born, 2013; Laventure et al., 2016). Moreover, it appears that sleep facilitates skill consolidation with a favourable molecular and cellular environment for plasticity (Abel et al., 2013), which is the key process of motor learning to improve performance and memory (Karni et al., 1995; Pascual-Leone et al., 1995; Rosenkranz et al., 2007).

Overall, our findings support the specific contribution of sleep in skill consolidation when the training is administrated in the evening. Thus, there may be a temporal window after training where sleep may provide consolidation benefits. A training session close to sleep could protect against memory deterioration induced by additional memories (retroactive interferences Brawn et al., 2010; Robertson, 2012). Accordingly, the morning group spending more time awake than the afternoon and evening group could be more exposed to interferences.

Another hypothesis for the deterioration of the morning group may be the involvement of different memory systems during the training.

The learning of a sequential finger tapping task requires both procedural (for skill) and declarative (for knowledge) memories (Breton and Robertson, 2014). While procedural memory appears to consolidate during wakefulness (Cohen et al., 2005; Robertson, 2009), the combination of procedural and declarative memories consolidate during sleep (Korman et al., 2003; Walker, 2003; Blischke et al., 2008; Doyon et al., 2009; Tucker et al., 2017). More interestingly, Brown et al. (2007) showed a conflict interaction between declarative and procedural memories, leading to a deterioration of performance. This conflict may be due to a common cerebral structure (medial temporal lobe) or a mediating structure (the dorsolateral prefrontal cortex) (Brown and Robertson, 2007; Robertson, 2012; Tunovic et al., 2014). In our study, the worst consolidation for the morning group could be due to the competition between both memories engaging in our finger tapping task. Further investigation is needed to provide a better understanding of the underlying mechanisms contributing to the time-of-day influence on consolidation processes.

Skill consolidation according to the time of day is a robust phenomenon and occurs regardless of age and chronotype. Indeed, better consolidation was found after evening training in both adolescent (Holz et al., 2012) and elderly (Korman et al., 2021) populations. Moreover, as our study included mainly intermediate-type chronotypes (see also Supplementary results 2 with only intermediate-type participants), Korman et al. (2021) showed the same consolidation result after evening training with the morning-type chronotype.

### Conclusion

The current study suggests that the time-of-day is an important factor that influences motor learning. Indeed, we showed that the consolidation fluctuates across the day, occurring better in the evening than in the morning. Even if new investigations are necessary to generalize these results and to better understand the underlying neural mechanisms, our findings may help to plan effective interventions in sports and rehabilitation.

## Data availability

The datasets generated during and analyzed during the current study are available from the corresponding author upon reasonable request.

## Author contributions

TC and PC designed the experiment; TC and RC recorded the data; TC and HPM analyzed the data; TC and RC developed figures; TC and PC wrote the manuscript; PC, RC, HPM, GJ and WO provided feedback on the manuscript; all co-authors read and approved the submitted version.

## References

Abel, T., Havekes, R., Saletin, J. M., and Walker, M. P. (2013). Sleep, Plasticity and Memory from Molecules to Whole-Brain Networks. Curr. Biol. 23, R774–R788. doi: 10.1016/j.cub.2013.07.025.

Albouy, G., Fogel, S., King, B. R., Laventure, S., Benali, H., Karni, A., et al. (2015). Maintaining vs. enhancing motor sequence memories: Respective roles of striatal and hippocampal systems. Neuroimage 108, 423–434. doi: 10.1016/j.neuroimage.2014.12.049.

Albouy, G., Sterpenich, V., Vandewalle, G., Darsaud, A., Gais, S., Rauchs, G., et al. (2013). Interaction between Hippocampal and Striatal Systems Predicts Subsequent Consolidation of Motor Sequence Memory. PLoS One 8, e59490. doi: 10.1371/journal.pone.0059490.

Atkinson, G., and Reilly, T. (1996). Circadian Variation in Sports Performance. Sport. Med. 21, 292–312. doi: 10.2165/00007256-199621040-00005.

Atkinson, G., and Speirs, L. (1998). Diurnal Variation in Tennis Service. Percept. Mot. Skills 86, 1335–1338. doi: 10.2466/pms.1998.86.3c.1335.

Blischke, K., Erlacher, D., Kresin, H., Brueckner, S., and Malangré, A. (2008). Benefits of Sleep in Motor Learning – Prospects and Limitations. J. Hum. Kinet. 20, 23–35. doi: 10.2478/v10078-008-0015-9.

Breton, J., and Robertson, E. M. (2014). Flipping the switch: mechanisms that regulate memory consolidation. Trends Cogn. Sci. 18, 629–634. doi: 10.1016/j.tics.2014.08.005.

Brown, R. M., and Robertson, E. M. (2007). Off-Line Processing: Reciprocal Interactions between Declarative and Procedural Memories. J. Neurosci. 27, 10468–10475. doi: 10.1523/JNEUROSCI.2799-07.2007.

Buysse, D. J., Reynolds, C. F., Monk, T. H., Berman, S. R., and Kupfer, D. J. (1989). The Pittsburgh sleep quality index: A new instrument for psychiatric practice and research. Psychiatry Res. 28, 193–213. doi: 10.1016/0165-1781(89)90047-4.

Cohen, D. A., Pascual-Leone, A., Press, D. Z., and Robertson, E. M. (2005). Off-line learning of motor skill memory: A double dissociation of goal and movement. Proc. Natl. Acad. Sci. 102, 18237–18241. doi: 10.1073/pnas.0506072102.

Coxon, J. P., Peat, N. M., and Byblow, W. D. (2014). Primary motor cortex disinhibition during motor skill learning. J. Neurophysiol. 112, 156–164. doi: 10.1152/JN.00893.2013/ASSET/IMAGES/LARGE/Z9K0131424780005.JPEG.

Dayan, E., and Cohen, L. G. (2011). Neuroplasticity Subserving Motor Skill Learning. Neuron 72, 443–454. doi: 10.1016/j.neuron.2011.10.008.

Debas, K., Carrier, J., Orban, P., Barakat, M., Lungu, O., Vandewalle, G., et al. (2010). Brain plasticity related to the consolidation of motor sequence learning and motor adaptation. Proc. Natl. Acad. Sci. 107, 17839–17844. doi: 10.1073/pnas.1013176107.

Dolfen, N., King, B. R., Schwabe, L., Swinnen, S., and Albouy, G. (2019). Glucocorticoid response to stress induction prior to learning is negatively related to subsequent motor memory consolidation. Neurobiol. Learn. Mem. 158, 32–41. doi: 10.1016/j.nlm.2019.01.009.

Doyon, J., and Benali, H. (2005). Reorganization and plasticity in the adult brain during learning of motor skills. Curr. Opin. Neurobiol. 15, 161–167. doi: 10.1016/j.conb.2005.03.004.

Doyon, J., Korman, M., Morin, A., Dostie, V., Tahar, A. H., Benali, H., et al. (2009). Contribution of night and day sleep vs. simple passage of time to the consolidation of motor sequence and visuomotor adaptation learning. Exp. Brain Res. 195, 15–26. doi: 10.1007/s00221-009-1748-y.

Driskell, J. E., Willis, R. P., and Copper, C. (1992). Effect of overlearning on retention. J. Appl. Psychol. 77, 615–622. doi: 10.1037/0021-9010.77.5.615.

Drust, B., Waterhouse, J., Atkinson, G., Edwards, B., and Reilly, T. (2005). Circadian Rhythms in Sports Performance—an Update. Chronobiol. Int. 22, 21–44. doi: 10.1081/CBI-200041039.

Edwards, B. J., Lindsay, K., and Waterhouse, J. (2005). Effect of time of day on the accuracy and consistency of the badminton serve. Ergonomics 48, 1488–1498. doi: 10.1080/00140130500100975.

Fischer, S., Hallschmid, M., Elsner, A. L., and Born, J. (2002). Sleep forms memory for finger skills. Proc. Natl. Acad. Sci. 99, 11987–11991. doi: 10.1073/pnas.182178199.

Fogel, S. M., Albouy, G., Vien, C., Popovicci, R., King, B. R., Hoge, R., et al. (2014). fMRI and sleep correlates of the age-related impairment in motor memory consolidation. Hum. Brain Mapp. 35, 3625–3645. doi: 10.1002/hbm.22426.

Guette, M., Gondin, J., and Martin, A. (2005). Time-of-Day Effect on the Torque and Neuromuscular Properties of Dominant and Non-Dominant Quadriceps Femoris. Chronobiol. Int. 22, 541–558. doi: 10.1081/CBI-200062407.

Gueugneau, N., and Papaxanthis, C. (2010). Time-of-day effects on the internal simulation of motor actions: psychophysical evidence from pointing movements with the dominant and non-dominant arm. Chronobiol. Int. 27, 620–639. doi: 10.3109/07420521003664205.

Gueugneau, N., Pozzo, T., Darlot, C., and Papaxanthis, C. (2017). Daily modulation of the speed–accuracy trade-off. Neuroscience 356, 142–150. doi: 10.1016/j.neuroscience.2017.04.043.

Hammerschmidt, D., and Wöllner, C. (2022). Spontaneous motor tempo over the course of a week: the role of the time of the day, chronotype, and arousal. Psychol. Res. 1, 1–12. doi: 10.1007/S00426-022-01646-2/FIGURES/2.

Holz, J., Piosczyk, H., Landmann, N., Feige, B., Spiegelhalder, K., Riemann, D., et al. (2012). The Timing of Learning before Night-Time Sleep Differentially Affects Declarative and Procedural Long-Term Memory Consolidation in Adolescents. PLoS One 7, e40963. doi: 10.1371/journal.pone.0040963.

Horne, J. A., and Ostberg, O. (1976). A self assessment questionnaire to determine Morningness Eveningness in human circadian rhythms. Int. J. Chronobiol. 4, 97–110.

Jasper, I., Häubler, A., Marquardt, C., and Hermsdörfer, J. (2009). Circadian rhythm in handwriting. J. Sleep Res. 18, 264–271. doi: 10.1111/j.1365-2869.2008.00727.x.

Karni, A., Meyer, G., Jezzard, P., Adams, M. M., Turner, R., and Ungerleider, L. G. (1995). Functional MRI evidence for adult motor cortex plasticity during motor skill learning. Nature 377, 155–8. doi: 10.1038/377155a0.

Keisler, A., Ashe, J., and Willingham, D. T. (2007). Time of day accounts for overnight improvement in sequence learning. Learn. Mem. 14, 669–672. doi: 10.1101/lm.751807.

King, B. R., Gann, M. A., Mantini, D., Doyon, J., and Albouy, G. (2021). Persistence of Hippocampal Multivoxel Patterns during Awake Rest after Motor Sequence Learning. bioRxiv, 1–21. doi: 10.1101/2021.06.29.450290.

Korman, M., Gal, C., Gabitov, E., and Karni, A. (2021). Better later: evening practice is advantageous for motor skill consolidation in the elderly. Learn. Mem. 28, 72–75. doi: 10.1101/lm.052522.120.

Korman, M., Raz, N., Flash, T., and Karni, A. (2003). Multiple shifts in the representation of a motor sequence during the acquisition of skilled performance. Proc. Natl. Acad. Sci. 100, 12492–12497. doi: 10.1073/pnas.2035019100.

Krakauer, J. W., Hadjiosif, A. M., Xu, J., Wong, A. L., and Haith, A. M. (2019). Motor learning. Compr. Physiol. 9, 613–663. doi: 10.1002/cphy.c170043.

Kuriyama, K. (2004). Sleep-dependent learning and motor-skill complexity. Learn. Mem. 11, 705–713. doi: 10.1101/lm.76304.

Kusumoto, H., Ta, C., Brown, S. M., and Mulcahey, M. K. (2021). Factors Contributing to Diurnal Variation in Athletic Performance and Methods to Reduce Within-Day Performance Variation: A Systematic Review. J. Strength Cond. Res. 35, S119–S135. doi: 10.1519/JSC.0000000000003758.

Kvint, S., Basiri-Tehrani, B., Pruski, A., Nia, J., Nemet, I., Lopresti, M., et al. (2011). Acquisition and retention of motor sequences: The effects of time of the day and sleep. Arch. Ital. Biol. 149, 303–312. doi: 10.4449/aib.v149i3.1244.

Lang, N., Rothkegel, H., Reiber, H., Hasan, A., Sueske, E., Tergau, F., et al. (2011). Circadian Modulation of GABA-Mediated Cortical Inhibition. Cereb. Cortex 21, 2299–2306. doi: 10.1093/cercor/bhr003.

Laventure, S., Fogel, S., Lungu, O., Albouy, G., Sévigny-Dupont, P., Vien, C., et al. (2016). NREM2 and Sleep Spindles Are Instrumental to the Consolidation of Motor Sequence Memories. PLOS Biol. 14, e1002429. doi: 10.1371/journal.pbio.1002429.

Lohse, K. R., Wadden, K., Boyd, L. A., and Hodges, N. J. (2014). Motor skill acquisition across short and long time scales: A meta-analysis of neuroimaging data. Neuropsychologia 59, 130–141. doi: 10.1016/J.NEUROPSYCHOLOGIA.2014.05.001.

McDonnell, M. N., Orekhov, Y., and Ziemann, U. (2007). Suppression of LTP-like plasticity in human motor cortex by the GABA B receptor agonist baclofen. Exp. Brain Res. 180, 181–186. doi: 10.1007/S00221-006-0849-0/FIGURES/2.

Morehead, J. R., Qasim, S. E., Crossley, M. J., and Ivry, R. (2015). Savings upon Re-Aiming in Visuomotor Adaptation. J. Neurosci. 35, 14386–14396. doi: 10.1523/JNEUROSCI.1046-15.2015.

Neville, K.-M., and Trempe, M. (2017). Serial practice impairs motor skill consolidation. Exp. Brain Res. 235, 2601–2613. doi: 10.1007/s00221-017-4992-6.

Oldfield, R. C. (1971). The assessment and analysis of handedness: The Edinburgh inventory. Neuropsychologia 9, 97–113. doi: 10.1016/0028-3932(71)90067-4.

Pan, S. C., and Rickard, T. C. (2015). Sleep and motor learning: Is there room for consolidation? Psychol. Bull. 141, 812–834. doi: 10.1037/bul0000009.

Pascual-Leone, A., Nguyet, D., Cohen, L. G., Brasil-Neto, J. P., Cammarota, A., and Hallett, M. (1995). Modulation of muscle responses evoked by transcranial magnetic stimulation during the acquisition of new fine motor skills. J. Neurophysiol. 74, 1037–1045. doi: 10.1152/jn.1995.74.3.1037.

Rasch, B., and Born, J. (2013). About Sleep’s Role in Memory. Physiol. Rev. 93, 681–766. doi: 10.1152/physrev.00032.2012.

Rickard, T. C., Cai, D. J., Rieth, C. A., Jones, J., and Ard, M. C. (2008). Sleep does not enhance motor sequence learning. J. Exp. Psychol. Learn. Mem. Cogn. 34, 834–842. doi: 10.1037/0278-7393.34.4.834.

Robertson, E. M. (2009). From Creation to Consolidation: A Novel Framework for Memory Processing. PLoS Biol. 7, e1000019. doi: 10.1371/journal.pbio.1000019.

Robertson, E. M. (2012). New Insights in Human Memory Interference and Consolidation. Curr. Biol. 22, R66–R71. doi: 10.1016/j.cub.2011.11.051.

Robertson, E. M., Pascual-Leone, A., and Miall, R. C. (2004a). Current concepts in procedural consolidation. Nat. Rev. Neurosci. 5, 576–582. doi: 10.1038/nrn1426.

Robertson, E. M., Pascual-Leone, A., and Press, D. Z. (2004b). Awareness Modifies the Skill-Learning Benefits of Sleep. Curr. Biol. 14, 208–212. doi: 10.1016/j.cub.2004.01.027.

Rosenkranz, K., Kacar, A., and Rothwell, J. C. (2007). Differential Modulation of Motor Cortical Plasticity and Excitability in Early and Late Phases of Human Motor Learning. J. Neurosci. 27, 12058–12066. doi: 10.1523/JNEUROSCI.2663-07.2007.

Ruffino, C., Truong, C., Dupont, W., Bouguila, F., Michel, C., Lebon, F., et al. (2021). Acquisition and consolidation processes following motor imagery practice. Sci. Rep. 11, 2295. doi: 10.1038/s41598-021-81994-y.

Sale, M. V., Ridding, M. C., and Nordstrom, M. A. (2008). Cortisol Inhibits Neuroplasticity Induction in Human Motor Cortex. J. Neurosci. 28, 8285–8293. doi: 10.1523/JNEUROSCI.1963-08.2008.

Sale, M. V., Ridding, M. C., and Nordstrom, M. A. (2013). Time of Day Does Not Modulate Improvements in Motor Performance following a Repetitive Ballistic Motor Training Task. Neural Plast. 2013, 1–9. doi: 10.1155/2013/396865.

Shea, J. B., and Morgan, R. L. (1979). Contextual interference effects on the acquisition, retention, and transfer of a motor skill. J. Exp. Psychol. Hum. Learn. Mem. 5, 179–187. doi: 10.1037/0278-7393.5.2.179.

Smith, M. A., Ghazizadeh, A., and Shadmehr, R. (2006). Interacting Adaptive Processes with Different Timescales Underlie Short-Term Motor Learning. PLoS Biol. 4, e179. doi: 10.1371/journal.pbio.0040179.

Spampinato, D., and Celnik, P. (2018). Deconstructing skill learning and its physiological mechanisms. Cortex 104, 90–102. doi: 10.1016/j.cortex.2018.03.017.

Spampinato, D., and Celnik, P. (2021). Multiple Motor Learning Processes in Humans: Defining Their Neurophysiological Bases. Neurosci. 27, 246–267. doi: 10.1177/1073858420939552.

Taylor, J. A., Krakauer, J. W., and Ivry, R. B. (2014). Explicit and Implicit Contributions to Learning in a Sensorimotor Adaptation Task. J. Neurosci. 34, 3023–3032. doi: 10.1523/JNEUROSCI.3619-13.2014.

Teo, W., Newton, M. J., and McGuigan, M. R. (2011). Circadian rhythms in exercise performance: implications for hormonal and muscular adaptation. J. Sports Sci. Med. 10, 600–6. Available at: http://www.jssm.org [Accessed March 24, 2021].

Truong, C., Hilt, P. M., Bouguila, F., Bove, M., Lebon, F., Papaxanthis, C., et al. (2022). Time-of-day effects on skill acquisition and consolidation after physical and mental practices. Sci. Rep. 12, 5933–5933. doi: 10.1038/S41598-022-09749-X.

Tucker, M. A., Morris, C. J., Morgan, A., Yang, J., Myers, S., Pierce, J. G., et al. (2017). The Relative Impact of Sleep and Circadian Drive on Motor Skill Acquisition and Memory Consolidation. Sleep 40. doi: 10.1093/sleep/zsx036.

Tunovic, S., Press, D. Z., and Robertson, E. M. (2014). A Physiological Signal That Prevents Motor Skill Improvements during Consolidation. J. Neurosci. 34, 5302–5310. doi: 10.1523/JNEUROSCI.3497-13.2014.

Verhoeven, F. M., and Newell, K. M. (2018). Unifying practice schedules in the timescales of motor learning and performance. Hum. Mov. Sci. 59, 153–169. doi: 10.1016/j.humov.2018.04.004.

Walker, M. P. (2003). Sleep and the Time Course of Motor Skill Learning. Learn. Mem. 10, 275–284. doi: 10.1101/lm.58503.

Walker, M. P., Brakefield, T., Allan Hobson, J., and Stickgold, R. (2003). Dissociable stages of human memory consolidation and reconsolidation. Nature 425, 616–620. doi: 10.1038/nature01930.

Walker, M. P., Brakefield, T., Morgan, A., Hobson, J. A., and Stickgold, R. (2002). Practice with sleep makes perfect: Sleep-dependent motor skill learning. Neuron 35, 205–211. doi: 10.1016/S0896-6273(02)00746-8.

Ziemann, U., Ilić, T. V, Iliać, T. V, Pauli, C., Meintzschel, F., and Ruge, D. (2004). Learning modifies subsequent induction of long-term potentiation-like and long-term depression-like plasticity in human motor cortex. J. Neurosci. 24, 1666–72. doi: 10.1523/JNEUROSCI.5016-03.2004.

