## Supplementary materials for "Time-of-day effects on motor learning"

### Supplementary results 1

#### Threshold values for the plateau performance

Since our threshold value of 0.001 was arbitrarily determined, we performed, as a control, the same analysis with a threshold value of 0.0002, 0.0003, 0.0004, and 0.0005. Table S1 indicates the mean (+SD) of the trial when groups reached the performance plateau according to the four different threshold values. One-way ANOVA did not found significant different between groups, whatever the threshold values chosen (with 0.0002: $F_{2,33} = 0.18$, $p = 0.83$, $\eta^2 = 0.01$; with 0.0003: $F_{2,33} = 0.11$, $p = 0.90$, $\eta^2 = 0.01$; with 0.0004: $F_{2,33} = 0.22$, $p = 0.80$, $\eta^2 = 0.01$; with 0.0005: $F_{2,33} = 0.05$, $p = 0.95$, $\eta^2 = 0.01$).

**Tableau S1.** Supplementary Result. The mean (+SD) of the first trial of the performance plateau according to the different threshold values for each group.

| | | First trial of the performance plateau | | | |
|---|---|---|---|---|---|
| **Groups** | | **0.0002** | **0.0003** | **0.0004** | **0.0005** |
| **$G10_{am}$** | Mean | 18 | 14 | 10 | 8 |
| | *SD* | *5* | *5* | *4* | *4* |
| **$G3_{pm}$** | Mean | 19 | 15 | 11 | 9 |
| | *SD* | *3* | *3* | *3* | *3* |
| **$G8_{pm}$** | Mean | 19 | 14 | 11 | 8 |
| | *SD* | *4* | *4* | *3* | *3* |

### Supplementary results 2

#### The G10am, G3pm, and G8pm groups without extreme and moderate chronotypes

Figure S1 shows the average values (SD) of skill after excluding from the statistical analysis 9 participants, 3 from each group with extreme and moderate chronotypes. rmANOVA revealed a significant interaction effect for skill ($F_{4,48} = 5.81$; $p < 0.001$, $\eta^2=0.33$). Post-hoc analysis did not show significant differences between groups in T1 (in all, $p > 0.8$). Skill significantly enhanced after training for all groups (T1 versus T2; in all, $p < 0.001$). The comparison of T1_T2 gain with the reference value *zero* (0) showed significant improvement in skill for all groups (in all, $t > 4.74$, $p < 0.01$); this improvement was similar between groups (one-way ANOVA: $F_{2,24} = 0.19$, $p = 0.82$, $\eta^2 = 0,02$). One day after training (T2 versus T3; *post hoc* analysis), we observed a deterioration in skill performance for the $G10_{pm}$ ($p = 0.009$), a stabilization for the $G3_{pm}$ ($p = 0.46$), and further improvement for the $G8_{pm}$ ($p = 0.02$). The T2_T3 gain comparison with the reference value zero (0) showed a deterioration of skill for the $G10_{am}$ ($t = -3.37$, $p = 0.01$; 8/9 participants decreased their performance), a stabilization for the $G3_{pm}$ ($t = 1.12$, $p = 0.30$; 6/9 participants increased their performance) and an improvement for the $G8_{pm}$ ($t = 2.70$, $p = 0.03$; 9/9 participants increased their performance). One-way ANOVA ($F_{2,24} = 9.25$, $p = 0.001$, $\eta^2 = 0,44$) showed that T2_T3 gain of the $G10_{am}$ significantly differed from that of the $G3_{pm}$ and $G8_{pm}$ (*post hoc* analysis: for both, $p < 0.01$). All groups acquired better skill performance one-day latter (T3) compared to their initial performance (T1 versus T3; *post hoc* analysis: in all, $p < 0.001$).

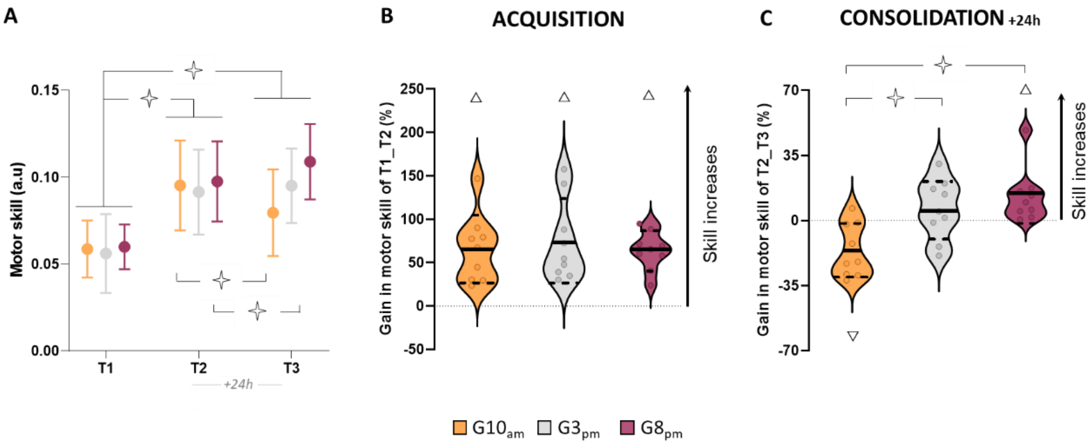


**Figure S1.** Supplementary Result. Skill performance for the G10am, G3pm, and G8pm groups without extreme and moderate chronotype. **(A)** Average values and standard deviations (+SD) of skill in T1, T2, and T3 for each group. **(B)** Violin plots for the percentage of acquisition gain in skill (T1_T2). Thick and thin horizontal lines mark mean and SD, respectively. Dots represent individual data per condition. **(C)** Trial-by-trial plotting of skill evaluation during the training and the corresponding power law functions. **(D)** Violin plots for the percentage of consolidation gain in skill (T2_T3) for each group. Thick and thin horizontal lines mark mean and SD, respectively. Dots represent individual data per condition. Open stars indicate significant differences between groups or sessions. Triangles indicate significant differences from the value *zero*.
